# Post-transcriptional control of EMT is coordinated through combinatorial targeting by multiple microRNAs

**DOI:** 10.1101/138024

**Authors:** Joseph Cursons, Katherine A Pillman, Kaitlin Scheer, Philip A Gregory, Momeneh Foroutan, Soroor Hediyeh Zadeh, John Toubia, Edmund J Crampin, Gregory J Goodall, Cameron P Bracken, Melissa J Davis

## Abstract

Epithelial-mesenchymal transition (EMT) is a process whereby cells undergo reversible phenotypic change, losing epithelial characteristics and acquiring mesenchymal attributes. While EMT underlies normal, physiological programs in embryonic tissue development and adult wound healing, it also contributes to cancer progression by facilitating metastasis and altering drug sensitivity. Using a cell model of EMT (human mammary epithelial (HMLE) cells), we show that miRNAs act as an additional regulatory layer over and above the activity of the transcription factors with which they are closely associated. In this context, miRNAs serve to both enhance expression changes for genes with EMT function, whilst simultaneously reducing transcriptional noise in non-EMT genes. We find that members of the polycistronic miR-200c~141 and miR-183~182 clusters (which are decreased during HMLE cell EMT and are associated with epithelial gene expression in breast cancer patients) co-regulate common targets and pathways to enforce an epithelial phenotype. We demonstrate their combinatorial effects are apparent much closer to endogenous expression levels (and orders of magnitude lower than used in most studies). Importantly, the low levels of combinatorial miRNAs that are required to exert biological function ameliorate the “off-target” effects on gene expression that are a characteristic of supra-physiologic miRNA manipulation. We argue that high levels of over-expression characteristic of many miRNA functional studies have led to an over-estimation of the effect of many miRNAs in EMT regulation, with over 130 individual miRNAs directly implicated as drivers of EMT. We propose that the functional effects of co-regulated miRNAs that we demonstrate here more-accurately reflects the endogenous post-transcriptional regulation of pathways, networks and processes, and illustrates that the post-transcriptional miRNA regulatory network is fundamentally cooperative.

## Introduction

Epithelial-Mesenchymal Transition (EMT) is the process by which epithelial cells alter their intercellular adhesion and acquire mesenchymal characteristics. These events involve molecular reprogramming of the cell, including loss or redistribution of epithelial-specific junctional proteins such as E-cadherin, rearrangement of the actin cytoskeleton, and induction of mesenchymal proteins such as vimentin, fibronectin and many others (Ye & Weinberg, 2015). In a mesenchymal state, cells possess an enhanced migratory ability which facilitates cell movement during developmental processes such as gastrulation and neural crest delamination and assists wound healing in the adult. There is also considerable evidence that EMT alters sensitivity to targeted drugs (Cursons et al., 2015; Foroutan, Cursons, Hediyeh-Zadeh, Thompson, & Davis, 2017) and chemotherapy (Arumugam et al., 2009; Fischer et al., 2015; Zheng et al., 2015). EMT can also promote metastasis, as it endows cancer cells with properties that facilitate detachment from the primary tumour, invasion through the basement membrane and entry into the circulation (Thompson & Haviv, 2011). EMT is a reversible processes, with a subsequent mesenchymal-epithelial transition (MET) facilitating colonisation and proliferation at secondary sites.

Many aspects of EMT regulation are well known: it is driven by coordinated changes in gene expression networks, controlled by both transcription factors (prominently from the ZEB, SNAIL and TWIST families) and miRNAs (such as the miR-200 family) which are linked in multiple co-regulatory relationships (Bracken, Scott, & Goodall, 2016; Friard, Re, Taverna, De Bortoli, & Cora, 2010; Gosline et al., 2016; Lu, Jolly, Levine, Onuchic, & Ben-Jacob, 2013; Re, Cora, Taverna, & Caselle, 2009). To date, more than 130 different miRNAs are reported to directly regulate EMT/MET (Table S1) with many of these studies heavily reliant upon the transient transfection of individual miRNAs to supraphysiological levels. However, as miRNAs work in a dose-dependent fashion, over-expression is likely to result in a multitude of off-target effects, where genes that would not be regulated by endogenous levels of the miRNA respond to supraphysiological abundances (Mayya & Duchaine, 2015; Witwer & Halushka, 2016). Understanding the roles of miRNAs is further complicated by the fact that EMT-promoting stimuli change the expression of dozens of miRNAs, but to levels that are typically lower than those achieved by transfection with miRNA mimics. This suggests that our interpretation of the role of individual miRNAs examined in isolation may not correspond to the endogenous functionality of multiple EMT-regulating miRNAs.

Individual transcripts can be regulated by multiple miRNAs (Wu et al., 2010) and single miRNAs can target multiple genes within the same biological pathways (Bracken et al., 2014; Bracken et al., 2016; Cai et al., 2015; Fujiwara et al., 2015; Jiang et al., 2016; Krishnan, Steptoe, Martin, Pattabiraman, et al., 2013; Krishnan, Steptoe, Martin, Wani, et al., 2013; Lal et al., 2011; C. W. Lin et al., 2013; Pellegrino et al., 2013; Peng et al., 2013; S. M. Tan et al., 2014; Wang et al., 2014). As one may expect, more target sites equate to more target repression (Doench, Petersen, & Sharp, 2003; Doench & Sharp, 2004; Krek et al., 2005; Nielsen et al., 2007; Pillai et al., 2005), particularly when target sites are optimally spaced (approximately 5-30nt apart) such that their co-operative activity is enhanced (Broderick, Salomon, Ryder, Aronin, & Zamore, 2011; Grimson et al., 2007; Hon & Zhang, 2007; Jopling, Schutz, & Sarnow, 2008; Rinck et al., 2013; Saetrom et al., 2007). Accordingly, functional cooperativity is observed between different miRNAs, particularly when they are polycistronic and share similar targets through related seed sites (Bracken et al., 2016; Gregory et al., 2008; Han et al., 2015; Khella et al., 2013; K. Y. Lin et al., 2016).

Despite an abundance of publications, significant questions remain regarding the role of miRNAs in EMT regulation: which miRNAs are truly capable of regulating the EMT/MET process *in vivo* and during cancer progression; to what extent do miRNAs regulate EMT at physiological concentrations, and; what are the endogenous roles of individual miRNAs given that EMT promoting stimuli drive co-ordinated changes in expression for dozens of miRNAs, both up-and down? Computational analyses have been used to predict cooperative miRNA targeting of individual mRNA transcripts (Schmitz et al., 2014). Similarly, computational studies have examined miRNA targeting of protein-protein interaction (PPI) networks (Zhu & Chen, 2014) and demonstrated a tendency of miRNAs to target protein-protein interaction network‘hub’ (highly connected) nodes (Liu, Guo, Pu, & Li, 2016). Many of these studies have stopped short of demonstrating the biological significance of these predicted regulatory interactions. Here, we explore the contribution of endogenous miRNAs to the regulation of EMT/MET using a model system that corresponds to the claudin-low, metaplastic subset of breast cancer which is associated with a poor clinical prognosis (Prat et al., 2010), and demonstrate the ability of selected miRNAs to control cellular phenotype in a combinatorial manner.

Using exon-intron split analysis (EISA) (Gaidatzis, Burger, Florescu, & Stadler, 2015), we demonstrate how miRNAs function as a secondary regulatory layer after transcription, amplifying transcriptional effects on EMT-associated processes (such as cell-adhesion and extracellular matrix organisation), whilst simultaneously buffering transcriptional effects of non-EMT genes. We also find that at physiologically relevant expression levels, EMT is regulated by multiple miRNAs acting in a combinatorial manner. In particular, we observe a dominant role for miR-200c-3p which is augmented by other miRNAs, including miR-141-3p, miR-182-5p and miR-183-5p. These miRNAs are not only co-regulated during TGFβ-induced EMT and across patient breast cancer samples, but computational analyses also suggest functional co-operation, co-targeting individual genes, and genes that are in close proximity within protein-protein interaction networks. Furthermore, as miRNA function is proportional to abundance, we find that transfection with multiple miRNAs at sub-nanomolar concentrations is sufficient to regulate EMT without off-target affects that are evident with transfection of miRNAs at high concentrations.

## Results

### The HMLE cell model of EMT shares a transcriptional profile with claudin-low metaplastic tumours

Exposing immortalised human mammary (HMLE) cells to TGFβ for 24 days drives the establishment of a stable sub-line (MesHMLE) with morphological (Fig. 1A), protein (Fig. 1B) and mRNA transcript (Fig. 1C) changes indicative of epithelial-mesenchymal transition (EMT). Subsequent ectopic expression of miR-200c in MesHMLE cells (MesHMLE+miR-200c) is sufficient to drive morphological changes (Fig. 1A) consistent with mesenchymal-epithelial transition (MET). However, molecular changes suggest MET is partial, as although there is increased *CDH1* (E-Cadherin) and reduced *ZEB1* expression at the protein (Fig. 1B) and mRNA level (Fig. S1A), the expression of vimentin remains high.

**Figure 1.**
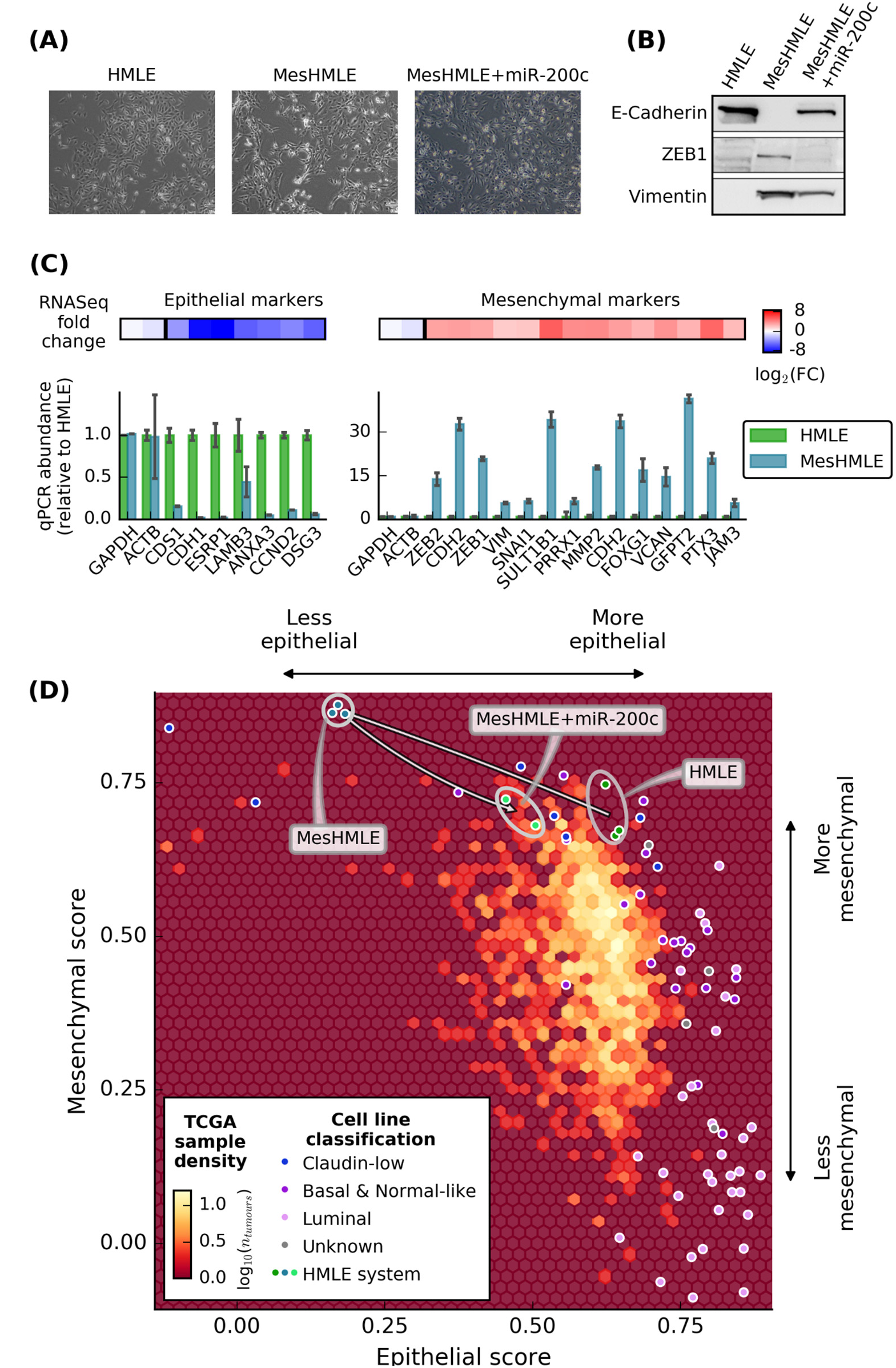
Characterising our human cell-line model of EMT. **(A)** The morphology of HMLE, MesHMLE (HMLE + 24 days TGFβ) and MesHMLE cells exposed to 20 nM of miR-200c mimic (MesHMLE + miR-200c). **(B)** Western Blots showing protein expression for selected epithelial (E-cadherin, encoded by *CDH1)* and mesenchymal (ZEB1, vimentin) markers. **(C)** Gene expression changes from RNA-Seq *(heatmap*, HMLE to MesHMLE) and qPCR transcript abundance data *(bar plots)* for epithelial *(left)* and mesenchymal *(right)* genes. *GAPDH* was used for qPCR normalisation and *ACTB* is an internal control. Error bars represent standard deviation. **(D)** A density plot of epithelial and mesenchymal score for all TCGA breast cancer samples *(hexbin)*, overlaid with a scatter plot of epithelial and mesenchymal scores for individual breast cancer cell lines.

We have previously characterised the relative epithelial and mesenchymal molecular phenotypes of cells on a two dimensional landscape (Foroutan et al., 2017), defined by separately scoring epithelial and mesenchymal gene sets (T. Z. Tan et al., 2014). In order to determine where our model system sits on this landscape, gene-set scores for the HMLE cell system were compared to TCGA breast cancer samples and a number of established breast cancer cell lines (Daemen et al., 2013; Heiser et al., 2012) (Fig. 1D). The TGFβ-driven transition between the HMLE and MesHMLE states is associated with a large reduction in the epithelial score and a smaller increase in the mesenchymal score of the cells (Fig 1D), while the following MesHMLE+miR-200c profile shows a similar mesenchymal score to the original HMLE cells, but only partially restores the epithelial score of the original HMLE cells. This is consistent with the morphology and markers (Fig 1 A-C) and supports the notion of partial MET driven by exogenous miR-200c exposure.

The MesHMLE cells also show a similar molecular phenotype to Basal B claudin-low cell lines (Blick et al., 2008) with very low epithelial scores (BT549 and HCC1395 cells; Fig. S1B). Further, MesHMLE cells also have epithelial and mesenchymal scores similar to a rare subset of TCGA breast cancer samples that typically have a metaplastic histological annotation (Table S2A). These cancers are characterised by lower expression of epithelial markers and higher expression of mesenchymal markers than in the remaining TCGA breast cancer data (Fig. S1C). The degree to which HMLE and MesHMLE cells are repositioned on the gene-set scoring axes suggests their epithelial/mesenchymal state is more heavily dependent upon changing expression of pro-epithelial genes rather than pro-mesenchymal ones.

### Post-transcriptional gene regulation simultaneously reinforces and buffers transcriptional changes during EMT

During EMT complex changes in gene expression are coordinated through both transcriptional and post-transcriptional regulation, primarily governed by their respective largest families of regulators: transcription factors (TFs) and miRNAs. To assess the relative contributions of these regulatory processes we used exon-intron split analysis (EISA) which maps reads to intronic and exonic regions (Gaidatzis et al., 2015). When an individual transcript is compared between two conditions, differences in the intronic reads (Δ*intron*) are indicative of altered transcription. Since exonic reads can come from mRNA transcripts before or after splicing, changes in the exonic reads (Δ*exon*) are influenced by both transcriptional and post-transcriptional effects; differences between exon and intron read changes (Δ*exon* – Δ*intron*) suggest altered post-transcriptional regulation.

The EISA metrics show that transcriptional and post-transcriptional regulation can either work in unison or opposition. Genes that are both transcriptionally and post-transcriptionally upregulated or downregulated we annotate as “co-ordinately increased” (CI) or “co-ordinately decreased” (CD) respectively. Genes where the transcriptional increase may be counteracted by post-transcriptional downregulation (or vice versa) we label as “increased, buffered” (IB) and “decreased, buffered” (DB) respectively (Fig. 2A; Table 1). As miRNAs regulate genes post-transcriptionally, when comparing HMLE and MesHMLE cells, one would anticipate that post-transcriptionally downregulated genes are targeted by miRNAs associated with a mesenchymal state. Alternatively, the loss of repression from miRNAs predominantly expressed in the epithelial state would drive post-transcriptional upregulation during EMT. The connectedness of the transcriptional and post-transcriptional arms may be further enhanced by miRNA-mediated regulation of the TFs themselves, with miRNA:TF feedback motifs a common feature of gene regulatory networks (Bracken et al., 2016; Lu et al., 2013).

**Figure 2.**
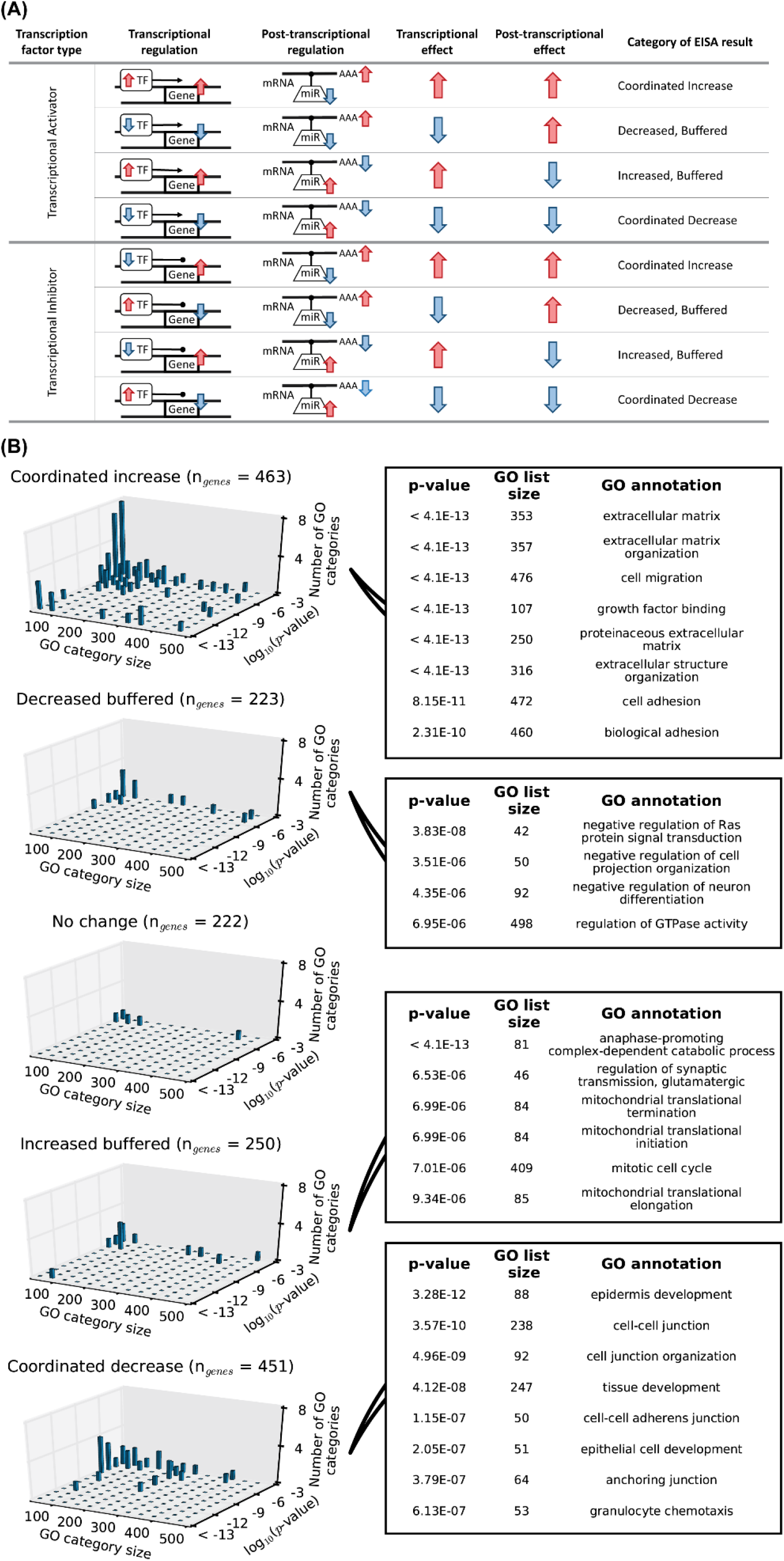
Post-transcriptional regulation can reinforce or buffer transcriptional changes associated with specific biological processes. **(A)** The categories into which genes were classified using exon-intron split analysis (EISA). **(B)** For each gene set the number of significant Gene Ontology (GO) annotations is shown, grouped across GO category list size and the degree of statistical support (log_10_(p-value)). The number of genes within each EISA partitioned set is given within the title of each sub-plot. For visualisation only significant (adjusted p-value <1x10^-3^) groups with a minimum GO list size of 40 are plotted *(3D histograms)*, while the largest and most significant GO categories are listed. More comprehensive lists are given in Table S4 *(Supplementary results)*.

**Table 1.** Gene sets were partitioned by changes to mRNA transcripts. Gene sets were partitioned for analysis using thresholds against mRNA transcript differential expression and exon-intron split analysis (EISA) metrics.

| Metric partitioned | Gene set label |  |  |  |  |
| --- | --- | --- | --- | --- | --- |
|  | Coordinated Increase | Decreased, Buffered | No change | Increased, Buffered | Coordinated Decrease |
| **Δintron** | Top 15% | Bottom 15% | -0.1 -> 0.1<br>(30<sup>th</sup> - 47<sup>th</sup> %ile) | Top 15% | Bottom 15% |
| **Δexon - Δintron** | Top 15% | Top 15% | -0.1 -> 0.1<br>(46<sup>th</sup> - 65<sup>th</sup> %ile) | Bottom 15% | Bottom 15% |
| **Differential transcript abundance** | - | - | -0.3 -> 0.31<br>(31<sup>st</sup> - 66<sup>th</sup> %ile) | - | - |

We found that genes subjected to co-ordinated up-or down-regulation were enriched for gene ontology terms that are of well-established importance during EMT (Fig. 2B). For example, genes annotated with terms including “extracellular matrix”, “cell migration” and “cell adhesion” were enriched among the co-ordinately upregulated gene set, whilst genes that were co-ordinately downregulated were involved in such processes as “cell-cell junction”, “cell junction organisation” and “epithelial cell development”. In contrast, genes whose expression was buffered by post-transcriptional regulation (where post-transcriptional control opposes transcriptional changes) had far fewer significantly enriched terms than genes that were co-ordinately regulated, despite similar numbers of genes in each group, indicating that the gene expression subject to post-transcriptional buffering included diverse genes with no strong functional similarity. Those few terms that were significant in the buffered sets had relatively small numbers of genes associated with them, and showed no obvious EMT associated functions. Transcription factors which were themselves part of the co-ordinately increased gene set include the transcriptional activators GLI2 and LEF1, the repressors ZEB1, ZEB2, TWIST1, TWIST2 and SMAD6, and the context-dependent RUNX2 (Table S3 *in Supplementary results).* Those within the co-ordinately decreased gene set include the activators EHF and ELF3, the repressors OVOL1 and OVOL2, and the context-dependent TFs GRHL2, GRHL3, KLF8 and SOX7 (Table S3).

These results indicate that concordant transcriptional and post-transcriptional regulation is a common cellular control mechanism during EMT and that post-transcriptional changes, largely mediated by miRNAs, reinforce both the up- and down-regulated transcription of EMT-associated genes. The lower number of enriched ontologies among the buffered gene sets is consistent with the notion that miRNAs simultaneously buffer transcriptional changes across a plethora of genes that are influenced by TF changes during EMT but do not necessarily have functions that are relevant to EMT processes.

### During EMT miRNAs co-regulate functionally-related transcripts

Post-transcriptional regulation appears to reinforce EMT transcriptional programs in our TGFβ-induced EMT model. Thus, we sought to characterise the roles of endogenous miRNAs in co-ordinating EMT gene regulation, beginning with small RNA-sequencing of HMLE and MesHMLE cells. We ranked miRNAs based on their relative abundance and their magnitude of differential expression (Fig. 3A; *see Methods/HMLE system miRNA ranking)*. In general, miRNAs that decreased in abundance have previously been described as pro-epithelial and associated with the induction of MET, while miRNAs that increased in abundance are generally pro-mesenchymal and associated with EMT (Table S1). An exception is miR-204, which has been described as pro-epithelial in a number of studies, but undergoes a large increase in abundance with the HMLE to MesHMLE transition (Fig. 3A).

**Figure 3.**
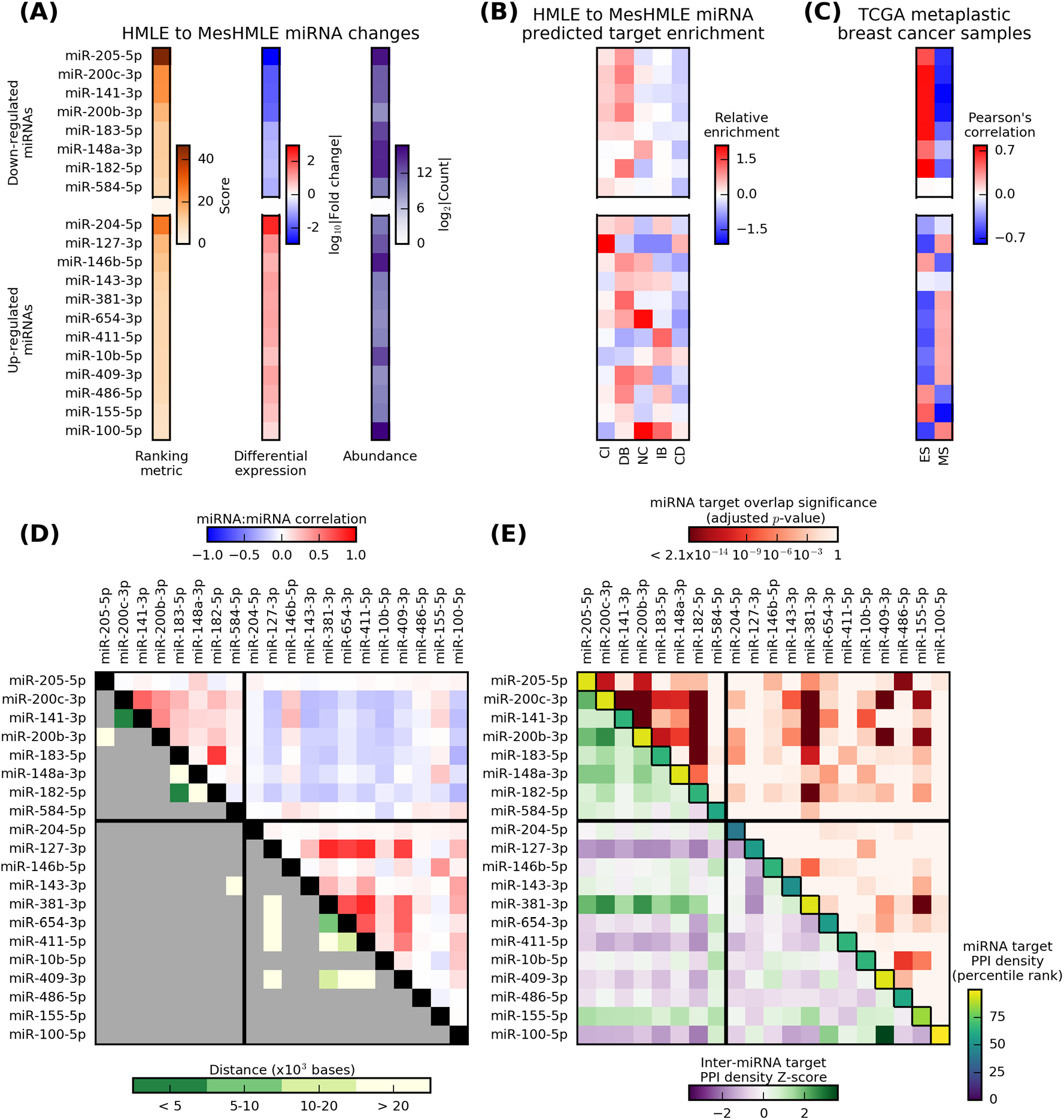
MiRNAs undergoing large changes in abundance during EMT in HMLE cells, show concordant changes within TCGA data and functional overlap in their predicted target genes. Full legend on previous page. **(A)** From the HMLE to MesHMLE transition, miRNAs were selected based upon their relative abundance (summed across both states) and differential expression *(details in Methods)*; the top 20 miRNAs are shown, split by the direction of their change in abundance. **(B)** The relative enrichment of high-confidence predicted targets (top 5% of TargetScan and DIANA-microT targets) for each miRNA within the EISA-partitioned gene sets (Fig. 2A). **(C)** Pearson's correlation between miRNA abundance and either epithelial score (ES) or mesenchymal score (MS) across metaplastic TCGA breast cancer samples (Table S2; n_tumours_=16). **(D)** Pearson's correlation between the abundance for each pair of miRNAs across all malignant TCGA breast cancer samples (at *top right)* as a measure of miRNA co-regulation across a larger set of tissue samples, together with the genomic distance between miRNAs located on the same chromosome (at *bottom left*; highlighting associations in abundance that can be attributed to polycistronic miRNA clustering). **(E)** Several metrics were applied to measure miRNA target set interactions (Fig. SM1 in *Supplementary methods)*. To quantify the enrichment of direct target overlap a x^2^-test was performed between each pair of miRNAs and adjusted p-values are shown (at *top right)*. To quantify relative dispersion of the predicted target set for an miRNA in the context of a protein-protein interaction (PPI) network, we calculated the density of interactions between predicted targets *(along diagonal)*. Similarly, to quantify the dispersion of targets for pairs of miRNAs, a bipartite graph was created from miRNA target set lists and the inter-miRNA target edge density was measured, and values are displayed as Z-score relative to all possible miRNA pairs (at *bottom left*).

To determine the contribution of miRNAs in controlling gene expression for our model we selected the 20 most abundant miRNAs (summed across both conditions; Fig. 3A) and calculated the relative enrichment of high-confidence predicted targets within each of the EISA-partitioned gene sets (Fig. 3B). For miRNAs downregulated during EMT our model of regulatory effects predicts that targets should be CI or DB, reflecting evidence of reduced post-transcriptional repression (Fig. 2A). With the exception of miR-148-3p, there is a consistent enrichment of targets within the co-ordinately increased (CI) and decreased buffered (DB) gene sets (Fig. 3B) as expected. Surprisingly, the inverse association for upregulated miRNAs was not present, and several miRNAs, including miR-127-3p, miR-381-3p, miR-409-3p and miR-486-5p, were found to have target transcripts enriched within the CI and DB sets.

As noted (Fig. 1D), the epithelial and mesenchymal transcript signature scores of MesHMLE cells are similar to TCGA breast cancer samples with characteristics of claudin-low breast cancer (Basal B cell lines and metaplastic tumours; Fig. S1 *in Supplementary Results)*, while HMLE cells occupy a region with both basal-like and metaplastic tumours (Fig. S3). Thus, we next investigated the candidate miRNAs identified from our HMLE/MesHMLE system within TCGA breast cancer data to determine whether their co-regulation was preserved across clinical samples. For most of the “pro-epithelial” miRNAs (miR-205, miR-200c, miR-141, miR-200b, miR-182 and miR-183) there was a particularly strong positive correlation with epithelial score and negative correlation with mesenchymal score, and this association was maintained but slightly weaker for miR-148a-3p (Fig. 3C). As expected, the “pro-mesenchymal” (EMT upregulated) miRNAs generally showed the opposite expression pattern across TCGA tumours, correlating negatively with epithelial score, although the strength of these associations tended to be lower than those of the top pro-epithelial miRNAs (Fig. 3C).

Given similarities between miRNA changes in the HMLE EMT model and miRNAs changing concordantly with epithelial and mesenchymal across the TCGA data, we further investigated potential miRNA co-regulation. Correlations were calculated between pairs of miRNAs across breast cancer samples, and as expected from associations with epithelial and mesenchymal score, many of these miRNAs showed evidence of cross-correlation (Fig. 3D; *top right).* It should be noted that the most positively correlated miRNA pairs, such as miR-200c/miR-141, miR-182/miR-183, and pairs of miR-381-3p/miR-654-3p/miR-411-5p, are all encoded at genomic loci within 20,000 bases (Fig. 3D; *bottom left)* and as such, these correlations largely reflect their polycistronic clustering. In addition however, several miRNAs that were not co-localised, such as miR-200c-3p and the miR-182-5p/183-5p cluster displayed a moderate positive association across breast cancers indicative of co-regulation.

We hypothesised that miRNAs are co-regulated in a manner which amplifies their function through cooperative targeting of functionally related transcripts. This might be evident as direct co-targeting of a transcript by co-expressed miRNAs, or by targeting of transcripts that encode proteins with functional relationships *(e.g*. direct protein-protein interactions; Fig. SM1 *in Supplementary Methods*). To assess miRNA direct co-targeting, we tested the null hypothesis that target overlap frequency is independent between miRNAs. Between many of the “pro-epithelial” miRNAs that were downregulated during EMT there was particularly strong evidence against this (Fig. 3E, *at top right*; adjusted p-value <1x10^-3^), suggesting enriched co-targeting, in particular between miR-200b-3p, miR-200c-3p, miR-141-3p and miR-182-5p (this is expected for miR-200b-3p and miR-200c-3p which share their seed sequence). It is interesting to note that amongst others, miR-381-3p and miR-409-3p had significant target set overlap with miR-200b-3p and miR-200c-3p (Fig. 3E, *at top right*). Accordingly, the apparent enrichment of miR-381-3p and miR-409-3p targets within the CI and DB gene sets (Fig. 3B) may reflect co-targeting between both “pro-epithelial” and “pro-mesenchymal” gene sets, which would suggest a complex network of balanced miRNA regulatory effects.

As another measure of potential functional overlap, we also quantified the connectivity of these high-confidence predicted miRNA target genes in the context of protein-protein interaction networks. For individual miRNAs, the PPI density of the target list (Fig. 3E; *diagonal elements)* reflects the relative dispersion of targets across targets with a functional interaction. We have previously reported that miR-200b targets multiple components of a highly connected cytoskeletal control network (Bracken et al., 2014), and this focussed targeting is reflected by the relatively high percentile rank of miR-200b-3p and miR-200c-3p. Amongst the other pro-epithelial miRNAs, miR-205-5p and miR-148a-3p show focussed targeting, while miR-183-5p and miR-182-5p appear to have slightly more dispersed targeting in the context of the protein interactome (Fig. 3E; *top left diagonal*). Finally, we also examined the PPI density of bipartite graphs induced from paired miRNA target lists, to quantify the degree of co-targeting around protein complexes and functionally interacting targets (Fig. 3E; *at bottom left*). In this context, there also appears to be evidence of relatively high co-targeting between the pro-epithelial miRNAs, suggesting that beyond direct co-targeting of shared target mRNA transcripts, there may also be cooperation between these miRNAs through the regulation of transcripts that encode functionally-interacting proteins.

Together, these results show highly co-regulated expression between the miR-200c/141 and miR-182/183 clusters (Fig. 3A, C &D), as well as strong evidence of cooperative function through both direct overlap of target transcripts, and through regulation of functionally interacting targets (Fig. 3E).

### Multiple miRNAs act in a combinatorial manner to promote MET

As demonstrated above, a number of pro-epithelial miRNAs from our EMT system show co-ordinated changes in breast cancer and are predicted to target functionally related genes, both through direct co-targeting of transcripts and in the context of protein-protein interaction networks. We tested the functional significance of these predicted co-regulators by re-expressing combinations of putatively pro-epithelial miRNAs in the MesHMLE cells and measuring the strength of the resulting MET.

Initially, we titrated the level miR-200c mimic to which MesHMLE cells were exposed, and as anticipated, there was a dose-dependent capacity to drive MET as indicated by the increased expression of epithelial marker genes (data not shown). We selected a low concentration of miRNA mimic (0.2nM, far lower than the 5-20uM typically used in miRNA studies) for the subsequent transfection of miRNAs into MesHMLE cells and observed at these very low transfection abundances, only miR-200c was capable of inducing epithelial marker genes *(CDH1, ESRP1*, and *CDS1)* on its own (Fig. 4A). Strikingly, although miR-141, miR-182 and miR-183 individually had no effect upon epithelial gene expression, they acted in a co-operative manner with miR-200c to strongly promote epithelial marker expression, indicating the importance of combinatorial miRNA activity at abundances at least an order of magnitude lower than those at which miRNAs are typically manipulated (Fig 4A). The relative contribution of miR-183 or miR-182 to epithelial gene expression is dependent upon the gene in question. Up-regulation of *CDH1* for example is dependent upon miR-200c and miR-182, whereas *CDS1* expression is more reliant upon miR-200c and miR-183 (Fig.4B). Furthermore, although miR-200c plays a driving role in the regulation of epithelial genes, it is the miR-183/-182 family, and not miR-200, that primarily suppresses vimentin (Fig.4C). This is likely to be via an indirect mechanism (much as the up-regulation of *CDH1* is mediated through miR-200-mediated repression of the ZEB1/ZEB2 transcription factors (Gregory et al., 2008) as neither the TargetScan nor DIANA-microT algorithms predict the direct targeting of vimentin by either miR-183 or miR-182. We observe a more prominent suppressive effect on the vimentin and ZEB1 proteins than the corresponding mRNAs by the miR-183/-182 and miR-200 families respectively (Fig.4 C & D) under these experimental conditions.

**Figure 4.**
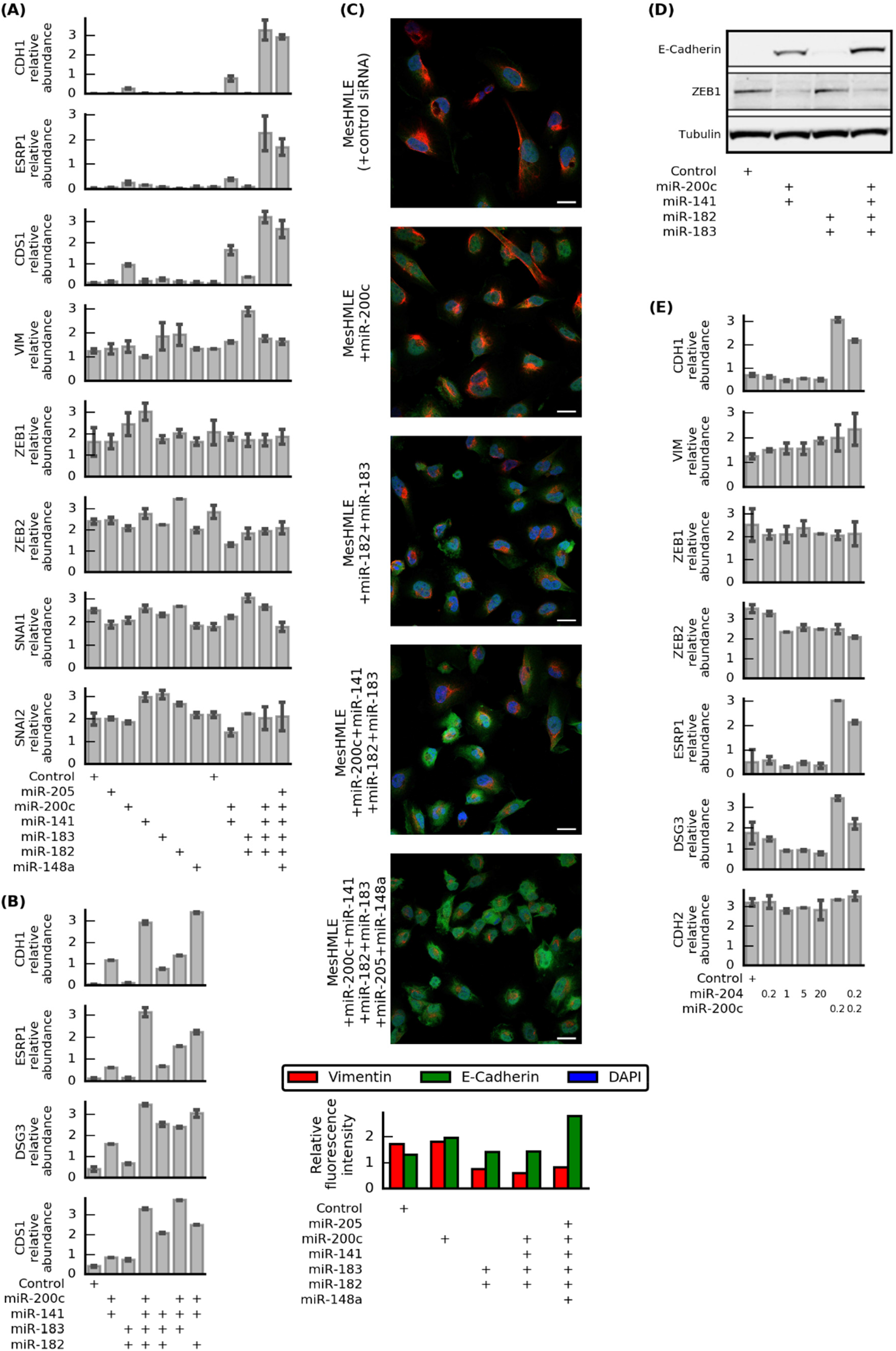
MiRNAs act in a combinatorial manner to promote an epithelial phenotype. **(A, B)** Marker gene expression in MesHMLE cells 3 days after transfection with 0.2 nM of the indicated miRNAs. **(C)** Expression of E‐‐cadherin and vimentin in MesHMLE cells detected by immunofluorescence 3 days after transfection with 0.2 nM of the indicated miRNAs. Scale bar represents 20 μm. Quantified fluorescence intensity *(inset)* was normalised against DAPI intensity. Scale bar represents 20 μm. **(D)** Expression of E-cadherin and ZEB1 by Western blotting following treatment with the specified combinations of miRNAs at 0.2 nM. **(E)** Expression of indicated markers genes with varying concentrations of miR-204 and/or 0.2 nM miR-200c.

As previously noted, a large increase in miR-204 abundance was observed with the HMLE to MesHMLE transition (Fig. 3A), contradicting a number of studies that have implicated miR-204 as pro-epithelial (Table S1). To better investigate its role in the MesHMLE system we expressed miR-204 alone at varying concentrations, and together at low levels with miR-200c (Fig. 4E). In this context, miR-204 alone appears to have little effect on *CDH1, CDH2* and *ESRP1* mRNA abundance, even with expression up to 20nM, although there may be minor effects upon ZEB2 and DSG3 abundance (consistent with an individually pro-epithelial role, at least when strongly over-expressed). When co-expressed with miR-200c, however, miR-204 antagonised the pro-epithelial effects of miR-200c, consistent with the notion that it may act to promote mesenchymal behaviour in this system. This emphasises the importance of testing miRNA function at physiological levels and in the context of other miRNAs.

### Ectopic expression of miRNAs targets genes that are not affected at endogenous miRNA levels

There is increasing evidence that the supraphysiological concentrations of miRNAs typical of most studies can result in overexpressed miRNAs targeting mRNA transcripts that are not regulated by that miRNA at endogenous expression levels (Mayya & Duchaine, 2015; Witwer & Halushka, 2016). As a consequence we have hypothesised, and others have recently shown, that many transcripts supposedly affected by miRNA in over-expression systems may not be biologically relevant (Pinzón et al., 2017). To investigate this in the context of our HMLE system, we compared EISA data for endogenous miRNA changes during the (TGFβ-driven) HMLE to MesHMLE transition against EISA data from the MesHMLE to MesHMLE+miR-200c transition, which involved exposure to ectopic miRNA at a concentration which was well in excess of physiological levels (Fig. 5).

**Figure 5.**
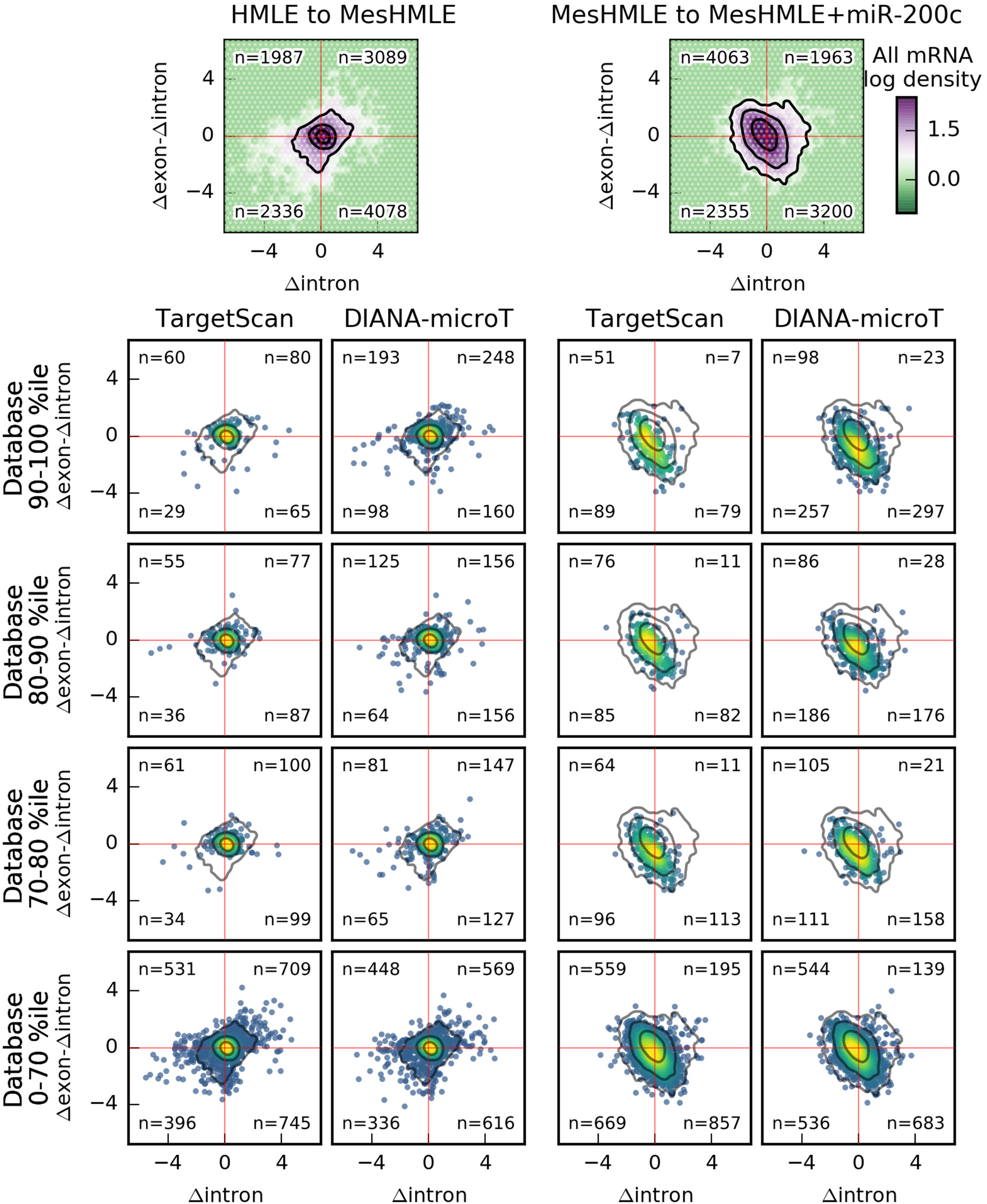
Differences in the transcriptional and post-transcriptional regulation of mRNA transcripts. The distribution of Aintron and Aexon-Aintron values are shown for mRNA transcripts during the HMLE to MesHMLE transition *(at left)* and the MesHMLE to MesHMLE+miR-200c transition *(at right)*. The density of all mRNA transcripts is shown (top row), and contour values are superimposed over the distribution of mRNA transcripts within the specified percentile range of database scores (at *left;* 10 percentile bins of *context+* for TargetScan and *miTG-score* for DIANA-microT). Red horizontal and vertical lines indicate the origin of each plot, and the number of mRNA transcripts within each resulting quadrant is shown. Note that there are differences in the number of miR targets across percentile bins of each database, and DIANA-microT in particular predicts a very large number of miR-200c-3p targets within the highest-scoring group (Fig. S4 *in Supplementary Results)*.

As noted above, non-zero values for hintron are indicative of altered transcription, while for Δexon-Δintron this is indicative of altered post-transcriptional regulation. For reference, the distribution for all mRNAs is shown *(top row)*, and density contours from these reference distributions are superimposed upon plots of mRNAs predicted to be miRNA targets with varying degrees of confidence *(as indicated by the percentile bins at left)*. The apparent rotation and skew of these distributions reflects the earlier observation that many genes experience transcriptional changes that are reinforced by concordant changes in post-transcriptional regulation.

For the HMLE to MesHMLE transition with an endogenous (approximately 80-fold) miR-200c decrease, there appears to be some enrichment of targets with reduced post-transcriptional control (Δexon-Δintron > 0) within the highest-confidence predicted targets *(i.e*. the 90^th^ – 100^th^ percentile of database metrics), however this enrichment is rapidly lost over lower scoring targets. Conversely, MesHMLE cells treated with a high concentration of miR-200c, there was evidence for increased post-transcriptional degradation across hundreds of predicted miR-200c targets, spanning a much greater range of prediction database scores (i.e. including more weakly predicted targets). These changes in the MesHMLE+miR-200c system imply that miR-200c expression directs the post-transcriptional degradation of thousands of transcripts. This highlights substantial differences between the activities of miRNAs at endogenous and exogenous levels of expression, where endogenously, miR-200c is a part-player in the EM or ME transition, whereas exogenously its overexpression alone is sufficient to drive the entire process. We strongly suspect the ability of over-expressed miRNAs to directly regulate thousands of targets (that would otherwise not be modulated at endogenous miRNA abundances), has led to a gross over-estimation for the role of individual miRNAs in EMT (Table S1).

### Combinations of miRNAs can induce MET with increased specificity

As discussed above, the overexpression of miR-200c drives the post-transcriptional degradation of thousands of predicted targets that do not appear to be under strong control by miR-200c during TGFβ-induced EMT. This suggests miR-200c over-expression may exert off-target effects that both wouldn't occur endogenously, and that may not occur if EMT is driven by combinations of miRNAs expressed at low (endogenous-like) levels. Consistent with previous observations, increasing levels of miR-200c result in increases in the expression of multiple epithelial genes in a dose-dependent manner *(CDH1, ESRP1, DSG3*, Fig. 6A) but not mesenchymal genes (with the prominent exception of *ZEB2*, Fig. 6B & C).

**Figure 6.**
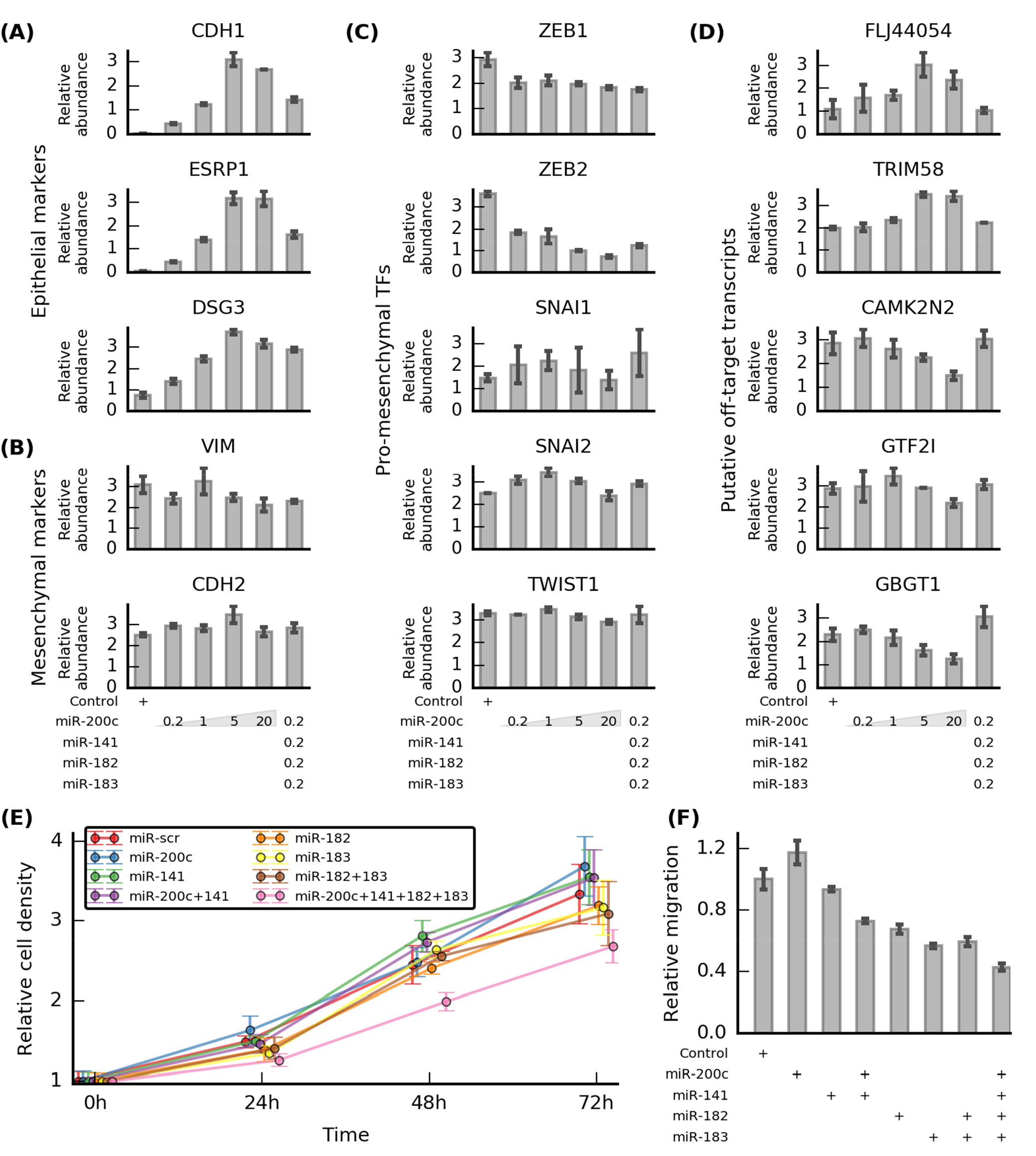
Transfection of sub-nanomolar concentrations of co-operative epithelial-enforcing miRNAs can drive MET with reduced “off-target” effects, reduced proliferation and reduced migration. The relative abundance of several transcripts was investigated after pro-epithelial miRNA transfection. Genes are divided into: **(A)** Known pro-epithelial (MET) markers, **(B)** known pro-mesenchymal EMT markers, **(C)** pro-mesenchymal EMT-promoting transcription factors, and **(D)** putative “off-target” genes that change in the MesHMLE to MesHMLE+miR-200 transition but not the TGFβ-induced HMLE to MesHMLE transition, are not previously associated with EMT, and do not have TargetScan-predicted sites for the miR-200 family, miR-182 or miR-183. Transfections (into MesHMLE cells) were performed in biological triplicate. qPCR was performed 72 hours post-transfection. Error bars represent standard deviation. The phenotypic effects of single miRNA and miRNA combination transfection was also assessed using **(E)** a cellular proliferation assay and **(F)** a trans-well migration assay. Assays were performed in triplicate and error bars represent standard deviation.

In order to ascertain “off-target” effects of over-expression, we identified genes that were: **(i)** altered in the RNA-seq data with ectopic miR-200c transfection (MesHMLE to MesHMLE+miR-200c) but not TGFβ induced EMT, (ii) not previously implicated in EMT, and (iii) not predicted targets of either the miR-200 or miR-182/183 families. We observed a number of these “off-target” genes were dose-dependently affected by miR-200c transfection, being either up-*(FLJ44056, TRIM58)* or down-regulated *(CAMK2N2, GTF2I, GBGT1)*. As shown previously (Fig. 4) and illustrated again here, members of the miR-200 and miR-182/183 families have combinatorial effects such that low-level co-transfection of these miRNAs has a co-operative effect on epithelial gene expression (Fig. 6A). Importantly however, despite still driving MET-like gene expression, these low level miRNA combinations no longer influenced off-target mRNAs (Fig. 6D).

Finally, we investigated the phenotype of MesHMLE cells following transfection of individual miRNAs or combinations of miRNAs using a proliferation and migration assay. Intriguingly, the relative density of cells transfected with miR-200c, miR-141, miR-182 and miR-183 was reduced compared to all other conditions at both the 48h and 72h time points (Fig. 6E). Furthermore, MesHMLE cells treated with miR-200c, miR-141, miR-182 and miR-183 in combination showed a reduced migratory capacity within a trans-well migration assay. These results show that low abundance pro-epithelial miRNAs can exert a combinatorial effect and induce a more robust MET than that which is achieved through treatment with extremely high concentrations of individual pro-epithelial miRNAs.

### Discussion

Post-transcriptional regulation by miRNAs is an active research field and there is growing interest in relating miRNAs to more general phenotypic programs such as EMT or cellular differentiation. EMT is a complex process where numerous stimuli can influence the regulatory activity of dozens of transcription factors to control expression across thousands of genes. The role of miRNAs in EMT is well-recognised, particularly for the miR-200 and miR-34 families which exist in reciprocal feedback loops with key TFs such as ZEB1/ZEB2 and SNAI1 to establish stable, self-sustaining epithelial or mesenchymal states (Lamouille, Xu, & Derynck, 2014; Saitoh, 2015; Tam & Weinberg, 2013). Over 130 miRNAs have been implicated in regulating EMT (Table S1), raising questions about the effects of common experimental manipulations *(e.g*. miRNA overexpression) and the extent to which these relationships are biologically meaningful, or even a reasonable reflection of endogenous function. As miRNAs can directly regulate a multitude of targets there has been a focus on characterising miRNA targets following manipulation; however it appears that the primary effects of miRNA perturbation are mediated by transcription factors (Gosline et al., 2016), and thus by solely characterising direct targets, the field has failed to capture the impact of miRNAs upon regulatory networks. Furthermore, it was recently reported that many predicted miRNA targets are functionally insensitive at endogenous miRNA levels and many miRNA target sites identified by computational algorithms may be conserved due to other evolutionary drivers (Pinzón et al., 2017). Motivated by these observations, we combined computational and experimental methods to identify multiple miRNAs with co-regulation and the potential for functional cooperation during EMT. The results presented here provide evidence for an alternative model of miRNA function, where small, simultaneous changes in the concentration of several molecules can exert strong functional effects through distributed targeting across molecular networks, with buffering of unrelated expression.

Working with an established model of TGFβ-induced EMT in breast cells (Conn et al., 2015; Lim et al., 2013) (Fig. 1) we used the exon-intron split analysis method (Gaidatzis et al., 2015) to separate transcriptional and post-transcriptional changes. Given the established role of alternative splicing in EMT (Lamouille et al., 2014) it is possible that differential exon usage affected the EISA metrics examined here, and some considerations for the EISA method are given within *Supplementary Methods*. A subsequent gene ontology enrichment analysis using EISA-partitioned data identified several EMT-associated transcript sets with increased abundance from promotion of transcription that was enhanced by a loss of repression from epithelial-promoting miRNAs, and similarly, a set of mesenchymal-associated miRNAs that supported the effects of repressive TFs which exerted their effects through EMT (Fig. 2A). It is interesting to note that a number of TFs within the co-ordinately increased and decreased gene sets are important regulators of EMT, in particular ZEB1/ZEB2 (Gregory et al., 2008; Lamouille et al., 2014; Perdigao-Henriques et al., 2016), RUNX2 (Chimge et al., 2011; Tan et al., 2016), and GRHL2 (Cieply, Farris, Denvir, Ford, & Frisch, 2013; Mooney et al., 2017), reflecting the high degree of interaction between transcriptional and post-transcriptional control in EMT. Given the relatively short sequence motifs through which TFs interact with target loci and their frequent role in controlling multiple processes it is likely that following TF perturbation a number of bystander genes may be effected. We observed several gene sets with evidence of post-transcriptional changes discordant to transcriptional changes, suggesting that beyond their role in reinforcing the activity of TFs, miRNAs have also evolved to buffer the effects of TF changes upon other genes. Accordingly, miRNAs may therefore, also play an important role in transcriptomic homeostasis extending beyond the well-studied miRNA:TF reciprocal inhibition motif (Bracken et al., 2016; Gregory et al., 2008; Tam & Weinberg, 2013) which plays a critical role in controlling network behaviour (Jolly et al., 2016; Lu et al., 2013).

Given the central role of EMT/MET in development it is essential that this program can be tightly regulated, and the observation above that over 130 miRNAs are proposed to individually modulate EMT (Table S1 in *Supplementary Results)* is difficult to reconcile with this notion. Many studies examine miRNA effects through exogenous expression, resulting in abundances that are hundreds of fold higher than natural endogenous levels detected by qPCR. This presents a particular challenge for miRNA research, as there is a gradient of targets: from those with strongly interactions through long (8nt) continuous complementary sequence between the miRNA and target with supplementary 3’ base pairing, through to transcripts with short, mismatched seed pairing sites that may not be regulated endogenously, but that may nevertheless become functional targets under conditions of high miRNA over-expression (Witwer & Halushka, 2016). We observe this clearly when comparing EISA profiles for HMLE cells that are exposed to TGF-β, where miR-200c-3p levels naturally change 80-fold, against the profiles of cells after exogenous miR-200c overexpression, where hundreds of weak target genes become miR-200c responsive (Fig. 5). We contend that non-physiological targeting by highly over-expressed miRNAs there is a confounding factor when interpreting the effects of individual miRNA manipulation to understand the normal functional role of that miRNA within cells. We see this even for miR-200c, perhaps the most well-known enforcer of an epithelial phenotype, which individually has minimal impact driving MET when expressed at low levels (Fig. 4 & 6). Instead, we have identified miRNAs that are co-regulated with miR-200c and predicted to co-target common protein complexes (Fig. 3), and we observe a dramatic additive effect when simultaneously modulating these miRNAs to promote MET (Figure 4) while minimising “off-target” effects that are seen when MET is induced by very high abundances of exogenous miR-200c (Fig. 6D). Induction of MET through simultaneous modulation of miR-200c, miR-141, miR-182, and miR-183 achieved a strong MET and had the strongest effect at reducing cellular migration (Fig. 6F), and intriguingly, caused the greatest reduction in cellular proliferation. In agreement with this, miR-200 family members have been shown to cooperate with the miR~183~96~182 cluster in prevent lung cancer invasion and metastasis (Kundu et al., 2016), although it should be noted that this previous work performed transfections with miRNA precursors at 50 or 100 nM, greatly exceeding the abundances used here.

Several of the miRNAs altered during the HMLE to MesHMLE transition (Fig. 3A) have been associated with mammary cell differentiation status, such that the downregulated miR-200 family members are expressed in luminal cells, while the upregulated miR-143, miR-100 and miR-204 have been linked to mammary stem cells/basal cells (Pal et al., 2015). Accordingly, it is tempting to speculate that the molecular changes we observed during this TGFβ-induced EMT correspond to a loss of cellular differentiation, while the phenotypic changes induced by our combinatorial miRNA treatment (Fig. 6E & F) reflect activation of a terminal differentiation program, rather than a pure induction of MET achieved through miR-200c alone. Indeed, the cooperation of these miRNAs in both lung (Kundu et al., 2016) and breast tissue suggests a more-general role in epithelialisation.

It appears that combinatorial effects of miRNAs are best characterised with regard to miRNAs co-regulated from polycistronic clusters. The miR-192-miR-194-miR-215 cluster, for example, can co-ordinately supress tumour progression in renal cell carcinoma (Khella et al., 2013). Similarly, each member of the let-7c-miR-99b-miR-125b cluster directly targets interleukin-6 receptor (IL-6R) and other components of the IL-6-signal transducer and activator of transcription 3 (STAT3) signalling pathway to decrease mammosphere growth, invasion and the metastatic spread of tumours in xenograft mouse models (K. Y. Lin et al., 2016). Targeted deletions within the miR-17~92 oncomiR cluster have also revealed functionally co-operative roles. Axial patterning for example was predominantly disrupted by the loss of miR-17, but further exacerbated by the additional loss of miR-18, whereas the loss of miR-19a and miR-19b impaired MYC-driven tumorigenesis. Cooperativity was further seen at the gene-expression level: the total number of genes disrupted by deleting the entire miR-17~92 cluster was far higher than the total number of genes disrupted by individual deletion of the corresponding individual miRNAs (Han et al., 2015). Furthermore, a recent report has shown that cooperative interaction with an adjacent target site can even confer function on a non-canonical‘seedless’ site that otherwise would fail to recruit RISC (Flamand, Gan, Mayya, Gunsalus, & Duchaine, 2017). Computational studies also suggest that clustered miRNAs co-regulate genes in shared protein-protein interaction networks and that the closer the proximity of proteins in the network, the more likely they are to be targeted by miRNAs from the same cluster (Yuan et al., 2009), but these predicted effects were not experimentally validated.

Beyond this, similar co-operativity has been reported for some non-clustered miRNAs. The mesenchymal-promoting SNAIL transcription factors for example are directly repressed by the miR-200, miR-203, and miR-34 families (Bracken, Khew-Goodall, & Goodall, 2015). The use of antisense inhibitors to miR-21, miR-23a and miR-27a together showed synergistic effects on reducing the proliferation of cells in culture and the growth of xenograft tumours in mice to a greater extent than the inhibition of single miRNAs alone (Frampton et al., 2014). Multiple miRNAs (miR-19a/b, miR-26a, miR-92) also co-target oncogenes including PTEN and BIM to reduce T-ALL cell viability (Mavrakis et al., 2011) and KLF4-mediated activation of both miR-21 and miR-206 can de-repress ERK signalling through co-targeting of RAS GTPase-activating protein 1 (RASA1) and sprouty-related EVH1 domain-containing protein 1 (SPRED1) (Sharma et al., 2014).

The work we present here not only provides another example of cooperativity between miRNAs, it demonstrates that at abundances nearer to endogenous levels of expression this cooperativity is essential for miRNAs to exert their biological effects. Further work is required to determine whether these miRNA targeting interaction effects are additive, or truly cooperative and synergistic. An important result was the observation that manipulating sets of miRNAs at low levels can reduce “off-target” effects that are seen with exposure to high abundances of individual miRNAs. This may expand therapeutic possibilities, as combinatorial targeting reduces the concentrations required for effective modulation of EMT programs.

In conclusion we have shown that combinatorial miRNA regulation provides a more efficacious approach to modulating cell phenotype with reduced off-target effects when compared to overexpression of individual miRNAs. The control of cell phenotype by miRNAs appears to be mediated through both the reinforcement of transcriptional changes and the buffering of changes to other important transcripts. Furthermore, the cooperative effects of simultaneously modulating several miRNAs at relatively low level abundances may be mediated through the co-targeting of protein signalling complex components. Using an established cell line model of EMT we have demonstrated that distributed, combinatorial targeting by multiple microRNAs can produce a more pronounced phenotypic effect than individual miRNAs used at higher concentrations. Together, these results provide support for the hypothesis that EMT is coordinated through distributed co-targeting by several microRNAs.

## Methods

### Computational analysis

The computational analysis was performed using python (v 3.6) with NumPy (Walt, Colbert, & Varoquaux, 2011), SciPy (Jones, Oliphant, & Peterson, 2014), and Matplotlib (Hunter, 2007). Public data were used in this analysis and details are given in Table SM1 (in *Supplementary Methods)*. Computational scripts are available under an MIT license, from:
http://github.com/DavisLaboratory/CombinatorialmiRNAs

### Cell lines, cell culture and miRNA transfection

Human Mammary Epithelial Cells (HMLE) cells were grown in HuMEC ready media (Gibco). A HMLE subline with a mesenchymal phenotype (MesHMLE) was achieved by culture in DMEM:F12 media (1:1) supplemented with 10 mg/ml insulin, 20 ng/ml EGF, 0.5 mg/ml hydrocortisone, and 5% fetal calf serum (FCS) and treatment with 2.5 ng/ml of TGFB1 (R & D) for 24 days. Where indicated, MesHMLE cells were transfected with miRNA mimics (GenePharma) using Lipofectamine RNAi-Max (Invitrogen) for 72h, with media replacement 12 hr post-transfection. For immunofluorescence, glass coverslips were inserted into each well and incubated in fibronectin solution (Roche: 10638039001; 25 μg per well at 50 μg/mL) at 37°C for 30 minutes. Excess fibronectin solution was removed using a pipette and cells were plated onto the coverslips.

### Isolation of RNA and Real-Time PCR

Total RNA was extracted using Trizol (Invitrogen) according to the manufacturer's instructions. Reverse transcription was performed using the SuperScript III cDNA synthesis kit (Invitrogen). Real-time PCR primers are listed in Table SM2 (in *Supplementary methods/PCR Primers)*, and values were quantified relative to the average values for GAPDH and HPRT.

### Western Blotting

Transfected MesHMLE cells were lysed in RIPA buffer (Abcam, ab156034) with protease inhibitor cocktail (Roche, 1183617001) and purified to produce whole cell extracts. 30 μg total protein was fractionated on 10% Bis-Tris Bolt gels (Life Technologies, NW00100B0X). A 100 mM Tris-Cl, 150mM NaCl, and 0.05% Tween (TNT) buffer at pH 7.5 was used. Proteins were transferred onto nitrocellulose membranes, blocked (5% skim milk powder in TNT) and probed in blocking buffer with the following primary antibodies: α-Tubulin (1:5000; Abcam, ab7291), ZEB1 (1:500; CST, D80D3), E-Cadherin (1:1000; BD-Transduction Laboratories, 610182) and Vimentin (1:1000; CST, 5741) overnight at 4°C. Blots were washed in TNT then probed with secondary antibodies (goat anti-rabbit HRP or goat anti-mouse HRP, Thermo Scientific; 31446 or 31460) and visualized on the Chemidoc Touch imaging system (BioRad) following application of Clarity Western ECL substrate (Biorad; 170-5060).

### Immunofluorescence

Following transfection, cells were washed with warm PBS and fixed in 4% paraformaldehyde (PFA) for 10 minutes. The cells were then permeabilized using wash buffer (0.1% Triton X-100/TBS) for 10 minutes, and blocked with 2% BSA in wash buffer for 1 hour at RT. Primary antibodies (1:500, BD-Transduction Laboratories; 610182) and Vimentin (1:100, CST; 5741) were diluted in block buffer and applied to the coverslips which were incubated overnight at 4°C. Following three 5 minute washes, the coverslips were incubated with goat anti-mouse AlexaFluor 488 (Thermo Fisher; A-11029) and anti-rabbit AlexaFluor 594 (Thermo Fisher; A-11037) for 1 hour at RT. After washing, coverslips were mounted using ProLong Gold antifade reagent with DAPI (Molecular Probes; P36935). Six images were taken per coverslip using the Zeiss LSM 700 confocal microscope. FIJI software was used to calculate the fluorescence intensity (integrated density) for each channel. These values were normalized to the DAPI stain and averaged to obtain the mean fluorescence intensity per condition.

### Cellular proliferation assay

Transfected MesHMLE cells were dissociated, counted, and replated into four 96 well plates in quadruplicate at a density of 2x10^3^ cells per well. Cells were left to settle at 37°C for 4 hours, after which plate 1 was removed (t=0). The media was aspirated and the plate was wrapped in parafilm and stored at -20°C. This was repeated for plates 2 (t=24h), 3 (t=48h) and 4 (t=72h). The CyQUANT Cell Proliferation Assay Kit (Invitrogen) was then used to quantify DNA content and thus, measure proliferation as per manufacturer instruction. CyQUANT GR dye was applied to each well and sample fluorescence immediately measured using the Fluostar, with filters set for excitation at 480nm and emission at 520nm. Data was blank corrected, averaged between technical replicates and normalised to gain the fold change from the t=0 time point for each transfection. Data is represented as mean fold change and standard deviation.

### Transwell migration assay

Transwell inserts were humidified with serum free media at 37°C for an hour. Meanwhile, transfected MesHMLE cells were dissociated, counted and resuspended in serum free media (DMEM + EGF + insulin + hydrocortisone). Media was aspirated from transwells, which were then placed above 600μL full media (5% added FCS). 2x10^5^ transfected cells were added to individual transwells in triplicate, and incubated at 37°C for 4 hours. Each transwell was rinsed in PBS, then fixed in buffered formalin overnight at 4°C. Following a PBS rinse, cells were removed from the inside membrane of the transwells using a cotton tip, and washed off with PBS. Transwells were incubated with 0.1% Triton X for 5 minutes to permeabilise the cells, which were subsequently rinsed with water and incubated with DAPI for 30 minutes at room temperature. The transwells were washed once more, the membranes dried and mounted onto microscope slides with fluorescent mounting medium (Dako). DAPI was imaged in 6 distinct locations per membrane. The number of cells per image was counted and the average per filter in turn averaged and normalised to the cell density of the original suspensions (determined by CyQUANT assay).

### RNA-sequencing

Poly(A) enriched mRNA (New England Biolabs, Ipswich, MA) from HMLE and MesHMLE cells was sequenced in triplicate biological replicates using the Illumina HiSeq 2500 platform with a stranded paired end protocol, and read length of 100. On average, 95 million raw paired reads were obtained for each sample (Table SM3 *in Supplementary methods)*.

For RNA-seq processing parameters please refer to the *Supplementary Methods*. Raw reads were adapter trimmed and filtered using cutadapt v1.8 (Martin, 2011), and for the resulting FASTQ files quality checks were performed with FastQC. Filtered reads were mapped against the human reference genome (hg19) using the TopHat (v. 2.0.9) spliced alignment algorithm (Kim et al., 2013) producing an average alignment rate of 91% (Table SM2 *in Supplementary methods)*. Alignments were visualised and interrogated using the Integrative Genomics Viewer v2.3.80 (Thorvaldsdottir, Robinson, & Mesirov, 2013). Cuffdiff (v. 2.1.1) (Trapnell et al., 2013) was used to calculate differential transcript abundance and estimate significance.

As detailed within the Supplementary Methods, the RNA-seq data were also processed in parallel. The trimmed read data (described above) were aligned to the hg19 reference genome using the R/Bioconductor package Rsubread (Chen, Lun, & Smyth, 2016). For the parallel exon-intron split analysis *(see below)* the intronic and exonic counts were summarised using featureCounts (Liao, Smyth, & Shi, 2014). For gene set scoring *(see below)* and batch analysis *(Supplementary methods)*, logCPM values were obtained using the R/Bioconductor package edgeR without TMM normalisation (Robinson, McCarthy, & Smyth, 2010).

### Exon-intron split analysis

The exon-intron split analysis (EISA) was performed essentially as described in Gaidatzis et al. (2015) (Gaidatzis et al., 2015). Custom python scripts were used for all steps. Briefly, using only non-overlapping genes and uniquely mapped reads, we quantified the number of reads in intronic and exonic regions in a strand-specific manner for all UCSC RefSeq mRNA transcripts from each gene. Read pairs were ‘exonic’ if the 5’ end of the first read was mapped within an exon of any UCSC transcript or ‘intronic’ if the first read mapped entirely within an intron and not within 10 base pairs of an exon. Genes with insufficient read coverage (less than 24 reads in either intron or exonic read count) were discarded and the data was log2 transformed after adding a pseudo-count of 8. For each sample, normalisation was performed separately for intronic and exonic counts by dividing by the total number of reads. To allow comparison between experiments, counts were then multiplied by a standardised ratio of coverage of 40 for exons and 2 for introns (the coverage, in millions of reads, for a representative polyA-selected sample). The analysis was performed for all possible combinations of pairs of samples where one sample was from each experimental group (e.g. Group A replicate 1 vs Group B replicate 1, A1 vs B2, A2 vs B1, A2 vs B2). For each pair of individual replicates, hexon and hintron were calculated as the difference in log2 read counts between experimental conditions. To summarise across replicates, the means of the hintron or hexon values were calculated; in text these are referred to as‘hintron’ and‘hexon'. The data were further filtered to ensure all of the examined genes were expressed (above the minimum coverage above) in all replicates in at least one experimental group.

As noted above, a parallel RNA-seq alignment and analysis was performed. Comparing the hintron and hexon values from this alternative mapping to the values obtained by the standard EISA analysis some quantitative differences were observed, although the results were largely consistent, particularly for the hexon data (Fig. SM5 in *Supplementary Methods)*.

### Gene set scoring

A full description of the gene set scoring approach is given in (Foroutan et al., 2017) and further technical details are given in the Supplementary Methods. For this analysis (Fig. 1D, Fig. S1B & Fig. S3) single-sample gene set enrichment analysis (Verhaak et al., 2013) (ssGSEA) was applied through the R/Bioconductor package GSVA (Hanzelmann, Castelo, & Guinney, 2013) to score each sample against epithelial and mesenchymal gene sets from Tan et al (T. Z. Tan et al., 2014). Samples examined with ssGSEA include the TCGA breast cancer data, and RNA-seq data from breast cancer cell lines (Daemen et al., 2013).

### TCGA Breast Cancer Data Analysis

TCGA Breast Cancer miRNA transcript data were downloaded on 21^st^ December 2016 using the genomic data commons client. Individual files were merged using python (v3.6) with the python data analysis library (pandas) (McKinney, 2010). The pre-merged mRNA transcript data were downloaded on 24^th^ February 2015 from the TCGA Data Portal. For each TCGA tumour sample the mRNA transcript data were used to calculate an epithelial and mesenchymal score as described above. The pandas corr function was used to calculate the Pearson's correlation between miRNA abundance and epithelial score or mesenchymal score, and between the abundance of miRNAs.

### Partitioning of gene sets by mRNA transcript differential expression

To partition gene sets which contain mRNAs that may be differentially regulated through alternative transcriptional and post-transcriptional processes (Fig. 2A), the top/bottom 15^th^ percentile of transcripts by Aintron and Aexon-Aintron were taken for regulated gene sets (Table 1). The gene set associated with no change was selected through Aintron, Aexon-Aintron and differential transcript abundances centred around zero, with the percentile values indicated (Table 1,‘No change’). The resulting number of transcripts within each gene set is shown on Fig. 2B.

### Gene Ontology analysis

Gene ontology (GO) (Ashburner et al., 2000; Gene Ontology, 2015) annotations were downloaded and using the underlying network structure *(i.e*. parent/child terms) GO membership was propagated up the annotation inheritance tree to improve coverage (Maetschke, Simonsen, Davis, & Ragan, 2012). To test the enrichment of membership for specific GO processes a x^2^-test was applied with expected values derived from genome-wide frequencies with one degree of freedom (gene is a member/not a member). GO annotations with less than 20 or more than 500 members were discarded. For multiple hypothesis testing a Bonferroni correction was applied using the number of remaining GO annotations (n=3691).

### HMLE system miRNA ranking

To identify relatively-high abundance miRNAs which underwent large differential changes during the HMLE to MesHMLE transition (Fig. 3A) we ranked miRNAs by the product of their summed abundance (x; across both HMLE and MesHMLE states) and the absolute value of the log2 fold change:

miRNA score = log_10_(x_HMLE_ + X_MesHMLE_) *log_2_|(x_hmle_/x_MesHMLE_)|

### Predicted miRNA targets

Predicted miRNA targets were taken from TargetScan (v7.1) (Friedman, Farh, Burge, & Bartel, 2009; Grimson et al., 2007; Lewis, Burge, & Bartel, 2005) for miRNA families, and from DIANA-microT CDS (Paraskevopoulou et al., 2013; Reczko, Maragkakis, Alexiou, Grosse, & Hatzigeorgiou, 2012) for individual miRNAs. To select for relatively high-confidence putative targets (Fig. S3 *in Supplementary Results)*, we took the high-confidence (top 5 percentile of) predictions from TargetScan (ranked by *context+)*, and DIANA-microT (ranked by *miTG-score)*.

There were some minor discrepancies where miRNAs mapped within the HMLE system data could not be directly mapped to the TargetScan database. In particular: hsa-miR-381 (data) was mapped to hsa-miR-381-3p; hsa-miR-411-5p was mapped to both hsa-miR-411-5p.1 and hsa-miR-411-5p.2; and hsa-miR-183-5p was mapped to both hsa-miR-183-5p.1 and hsa-miR-183-5p.2. It should be noted, however, that at least in the case of miR-183-5p, different isomiRs have shown enrichment in triple negative breast cancer which is dependent upon patient race (Telonis, Loher, Jing, Londin, & Rigoutsos, 2015), and these effects are not considered here.

### MiRNA targeting analysis

To examine the enrichment of high confidence miRNA targets within the EISA-filtered gene sets (Fig. 3B), the relative fraction of transcripts within each EISA-partitioned group (Fig. 2A) was determined, and for each miRNA this was used to calculate the number of targets expected within each EISA-filtered set. The relative enrichment was calculated as: *(n_observed_* - *n_expected_)* / *n_expected_*.

A schematic illustration of target overlap quantification (Fig. 3E) is shown in Figure SM1 *(Supplementary Methods/Target set overlap and protein-interaction connectivity for* miRNAs), together with an expanded description of the method. Briefly, to examine the significance of direct miRNA target set overlap (Fig. 3E, *at top right)*, the frequency of targets for a miRNA was used to calculate an expected frequency of shared targets when compared to another miRNA, under the assumption of independence between miRNA target sets (Fig. SM1A). A χ^2^-statistic was calculated using the observed and numbers of shared targets, and a χ^2^-test was applied with 1 degree of freedom to estimate a p-value for evidence against the null hypothesis that miRNA target sets are independent. A Bonferroni correction was applied, and adjusted p-values were calculated by multiplying the estimated p-value by the total number of miRNA pairs: *(n_miRNAs_*(n_miRNAs_* - 1)) / 2.

Next, a protein-protein interaction (PPI) network was constructed by combining experimentally observed interactions from BioPlex (Huttlin et al., 2015), PINA v2.0 (Cowley et al., 2012) and InnateDB (Breuer et al., 2013). For each individual miRNA (Fig. 3E, *along diagonal)* we calculated the density of the PPI subgraph generated from high confidence predicted miRNA targets, and by comparing to the distribution across all miRNAs the percentile rank was determined (Fig. SM1B). For each pair of miRNAs (Fig. 3E, *bottom left)* we used the target lists to construct a bipartite graph, and calculated the resulting graph density. The distribution of density values was Gaussian and approximately Normal, thus density values (ρ) were converted to a Z-score using the sample mean (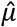)and standard deviation 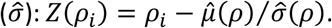 As illustrated (Fig. SM1C), this approach ignores nodes (mRNA transcripts) targeted by both miRNAs and edges (protein interactions) between targets of an individual miRNA; however both of these features will be captured by the preceding metrics (Fig. SM1A & B).

## Data Access

At submission the RNA-seq data used in this study are under preparation for upload to the European Nucleotide Archive. Please contact the authors for further information, or check for a subsequent version of this manuscript. Public data used in this report are listed within Table SM1.

## Acknowledgements

The authors would like to acknowledge: Prof. E Thompson from Queensland University of Technology for his advice on breast cancer subtypes; Daniel Esposito from the Walter and Eliza Hall Institute for the combined the PPI databases. Results published here are, in part, based upon data generated by the TCGA Research Network: http://cancergenome.nih.gov/.

This work was supported in part by: National Breast Cancer Foundation funding of the EMPathy Breast Cancer Network (CG-10-04), a National Collaborative Research Program; the Australian Research Council Centre of Excellence in Convergent Bio-Nano Science and Technology (project number CE140100036); and Australian National Health and Medical Research Council project grants GNT1068773 (GJG and PAG), GNT1034633 and GNT1069128 (GJG and CPB).

MF supported by Melbourne International Fee Remission Scholarship (MIFRS) and Melbourne International Research Scholarship (MIRS). PAG supported by Cancer Council SA Beat Cancer Fellowship. GJG supported by fellowships from the Australian National Health and Medical Research Council (GNT1026191 and GNT 1118170). CPB supported by Florey Fellowship from the Royal Adelaide Hospital Research Foundation. MJD funded by National Breast Cancer Foundation ECF-14-043.

## Author Contributions

JC, PAG, EJC, GJG, CPB and MJD designed the study; KS, PAG and CPB generated data; JC, KAP, KS, MF, SHZ, JT, CPB and MJD performed data analysis; and JC, CPB and MJD wrote the manuscript with input from all other authors.

